# Modular mRNA platform for *in vivo* antigen-specific immune tolerance

**DOI:** 10.64898/2026.05.18.725807

**Authors:** Shota Imai, Naoto Nishida, Ryo Maeno, Ayano Ikebuchi, Hitoshi Sasatsuki, Risa Saito, Kazutaka Matoba, Sadahiro Iwabuchi, Tomohiro Iba, Hisamichi Naito, Kanto Nagamori, Toan Van Le, Iriya Fujitsuka, Makie Ueda, Rikinari Hanayama, Tomoyoshi Yamano

## Abstract

Immune-mediated diseases such as autoimmunity and allergy arise from dysregulated antigen-specific responses, yet current therapies largely rely on broad immunosuppression and rarely re-establish durable, antigen-selective tolerance. Here we introduce Tol-mRNA, a modular tolerogenic mRNA platform designed to deliver multiple immune-regulatory signals *in vivo* to program antigen-specific tolerance. Tol-mRNA co-encodes an antigen together with two complementary regulatory cues, PD-L1 and TGF-β, enabling coordinated antigen presentation and tolerogenic signaling that robustly induces antigen-specific Tregs and suppresses pathogenic effector responses.

In murine disease models, Tol-mRNA conferred therapeutic benefit across distinct immune pathologies. In experimental autoimmune encephalomyelitis, Tol-mRNA ameliorated clinical disease and reduced inflammatory immune activation. In an ovalbumin-driven food allergy model, Tol-mRNA prevented allergic symptoms and dampened type 2 inflammation. Extending these findings to humans, Tol-mRNA demonstrated potent, antigen-dependent immunosuppressive activity and a strong capacity to promote antigen-specific Treg induction in human immune settings. Collectively, these results establish Tol-mRNA as a scalable and versatile approach to reprogram antigen-specific immunity and support its development as a next-generation therapeutic modality for immune-mediated diseases.

## Introduction

Autoimmune and allergic diseases arise when immune responses against self or environmental antigens become dysregulated, reflecting a fundamental breakdown of immune tolerance^1–4^. Autoimmune disorders are driven by pathogenic T and B cells that attack host tissues^5^, whereas allergic diseases are characterized by exaggerated type 2 immunity toward innocuous antigens such as food proteins, pollen, or dust mites^6^. Despite their distinct effector mechanisms, both disease classes share a common underlying biology: the failure to establish or maintain antigen-specific tolerance.

Current therapies, including systemic immunosuppressants, corticosteroids, and cytokine-targeting biologics, can ameliorate symptoms but do not restore immune homeostasis and often impair protective immunity^2^. As a result, patients frequently require long-term treatment without achieving durable disease remission^7^.

Regulatory T cells (Tregs) play an essential role in sustaining peripheral tolerance, and defects in Treg number or function are implicated in numerous autoimmune and allergic disorders^8–10^. However, adoptive transfer of polyclonal Tregs lacks antigen specificity, raising safety concerns such as suppression of beneficial immune responses^10^. These limitations underscore the need for antigen-specific strategies that reconstitute tolerance in a precise and physiological manner. In natural settings, peripherally induced Tregs (pTregs) are generated when naïve CD4⁺ T cells encounter antigen in the presence of a coordinated set of tolerogenic cues^11,12^. These include TGF-β–mediated transcriptional programming, co-inhibitory signals such as PD-L1, and specialized tolerogenic dendritic-cell (DCs). This process requires the simultaneous integration of antigen presentation and multiple regulatory signals, forming a tolerogenic microenvironment that is challenging to reconstruct synthetically. Recent mRNA-based approaches have demonstrated the feasibility of inducing antigen-specific tolerance by delivering disease-relevant antigens *in vivo*^13,14^. However, existing strategies typically provide antigen alone or pair it with a single immunoregulatory factor, and thus fail to reproduce the complexity of natural tolerogenic signaling required for stable and robust Treg induction.

To overcome these limitations, we developed Tol-mRNA, a modular tri-component tolerogenic mRNA platform designed to synthetically reconstitute the integrated signaling environment necessary for antigen-specific Treg differentiation. Tol-mRNA consists of three coordinated mRNA components: an antigen-encoding mRNA, a PD-L1 mRNA, and a TGF-β mRNA. Its modular design allows flexible replacement of antigen sequences while relying on fixed PD-L1 and TGF-β regulatory mRNAs, providing broad applicability across immune-mediated diseases.

We hypothesized that the coordinated *in vivo* delivery of these three tolerogenic signals would more faithfully mimic physiological Treg induction and provide a generalizable strategy for establishing antigen-specific immune tolerance. In this study, we describe the design and functional validation of Tol-mRNA and evaluate its potential as a broadly applicable platform for treating autoimmune, allergic, and other tolerance-related disorders.

## Results

### The Tol-mRNA induces antigen-specific regulatory T cells *in vivo*

Tolerogenic DC subsets are known to play a central role in the induction and maintenance of peripheral Tregs. These cells promote the differentiation of Tregs through controlled antigen presentation, low expression of costimulatory molecules, high expression of PD-L1, and the production of immunoregulatory cytokines such as TGF-β^11,12^. To efficiently induce antigen-specific Tregs *in vivo*, we designed a Tol-mRNA refers to a three-component mRNA formulation consisting of antigen-encoding mRNA, PD-L1 mRNA, and TGF-β mRNA.

We first evaluated the ability of the Tol-mRNA to induce antigen-specific Tregs against myelin oligodendrocyte glycoprotein (MOG), a well-established autoantigen in experimental autoimmune encephalomyelitis (EAE). The combination of antigen peptide, TGF-β, and PD-L1 induced significantly higher frequencies of antigen-specific Tregs compared with antigen peptide alone (Fig. 1a). In addition to MOG, the Tol-mRNA induced antigen-specific Tregs against multiple antigens, including ovalbumin (OVA) as well as type 1 diabetes–associated antigens insulin β and HIP2.5 (Fig. 1b). In contrast, a combination of peptide, CD80, and IL-2 promoted expansion of Foxp3 negative antigen-specific T cells (Extended Data Fig. 1).

**Figure 1.**
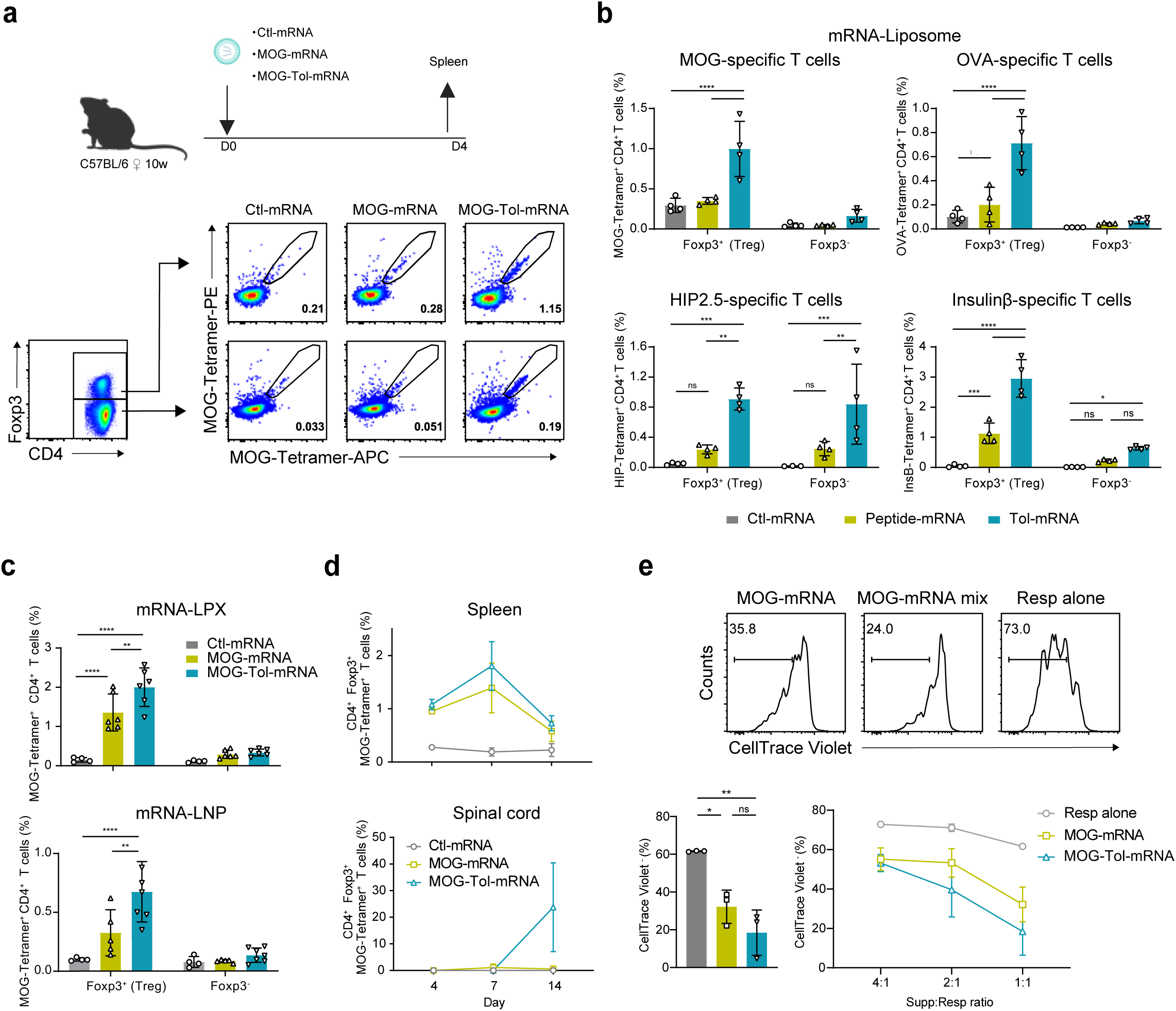
Induction of antigen-specific Tregs by Tol-mRNA. (a and b) Frequencies of antigen-specific CD4⁺ T cells within Foxp3⁺ Tregs or Foxp3⁻ conventional T cells were analyzed in the spleens of mRNA-treated mice. In C57BL/6 mice, MOG– and OVA–specific CD4⁺ T cells were evaluated, whereas in NOD mice, insulin-β– and HIP2.5–specific CD4⁺ T cells were assessed. (c) Frequency of splenic MOG–specific CD4^+^ T cells in Foxp3^+^ Tregs or Foxp3^-^ T cells after mRNA-LPX or mRNA-LNP injection. (d) Proportion of MOG–specific Tregs in the spleen and spinal cord on days 4, 7, and 14 after mRNA administration. (e) MOG-specific CD4⁺ T cells were enriched after mRNA treatment. These cells were co-cultured with CTV-labeled responder 2D2 T cells (CD45.1/2) in the presence of MOG peptide–loaded splenocytes (CD45.1) as stimulator cells. Proliferation of responder T cells was assessed by CTV dilution.

Lipoplex (LPX) and lipid nanoparticle (LNP) systems are among the most established platforms for mRNA delivery^15,16^. Both mRNA-LPX and mRNA-LNP induced antigen-specific Tregs more efficiently than antigen peptide alone; however, mRNA-LPX showed the highest induction efficiency and was therefore selected as the lipid carrier for subsequent experiments (Fig. 1c).

We next examined the *in vivo* persistence of induced antigen-specific Tregs. In both the peptide mRNA and Tol-mRNA, antigen-specific Tregs expanded in the spleen up to day 7 and declined by day 14. The expanded antigen-specific Tregs highly expressed Treg-associated molecules (Extended Data Fig. 2). Notably, antigen-specific Tregs induced by the Tol-mRNA, but not those induced by MOG-mRNA, were detected in the spinal cord (Fig. 1d). In *in vitro* suppression assays, antigen-specific Tregs induced by the Tol-mRNA exhibited superior suppressive activity compared with those induced by peptide mRNA alone (Fig. 1e).

Collectively, these results demonstrate that the Tol-mRNA efficiently induces durable and functionally suppressive antigen-specific Tregs *in vivo*.

### MOG-Tol-mRNA prevents disease progression in EAE

We evaluated whether induced antigen-specific Tregs could ameliorate autoimmune disease symptoms. To test this, we employed the EAE model ^17^. The MOG-Tol-mRNA showed significantly reduced disease severity compared to the Ctl-mRNA and MOG-mRNA groups (Fig. 2a, Extended Data Fig. 3a). In addition, body weight loss in EAE mice was significantly suppressed (Extended Data Fig. 3b). Histological analyses revealed demyelination and prominent inflammatory cell infiltration in the spinal cords of mice treated with Ctl-mRNA or MOG-mRNA, whereas these pathological features were substantially improved in the MOG-Tol-mRNA-treated mice (Fig. 2b, Extended Data Fig. 3c). Infiltration of CD4⁺ T cells, as well as MOG-specific Th1 and Th17 cells in the spinal cord, was reduced by the MOG-Tol-mRNA (Fig. 2c, d). The number of microglia in the spinal cord was highest in the Tol-mRNA treated group, and high infiltration of macrophages, DCs, and moDCs was observed in the control groups (Fig. 2e). These cells maintained high MHCII and CD86 expression, which may have been associated with disease exacerbation (Extended Data Fig. 3d). A similar trend to that observed for spinal cord–infiltrating T cells was also seen in the spleen. MOG-specific Th1 and Th17 cells in the spleen were similarly reduced by the MOG-Tol-mRNA (Fig. 2f). Polyclonal T cells within the spleen showed a decreasing trend in the mRNA therapy group, but no differences were observed in other immune cells (Extended Data Fig. 3e). Furthermore, cytokines associated with disease progression in both the spinal cord and spleen were significantly decreased in the MOG-Tol-mRNA (Extended Data Fig. 4).

**Figure 2.**
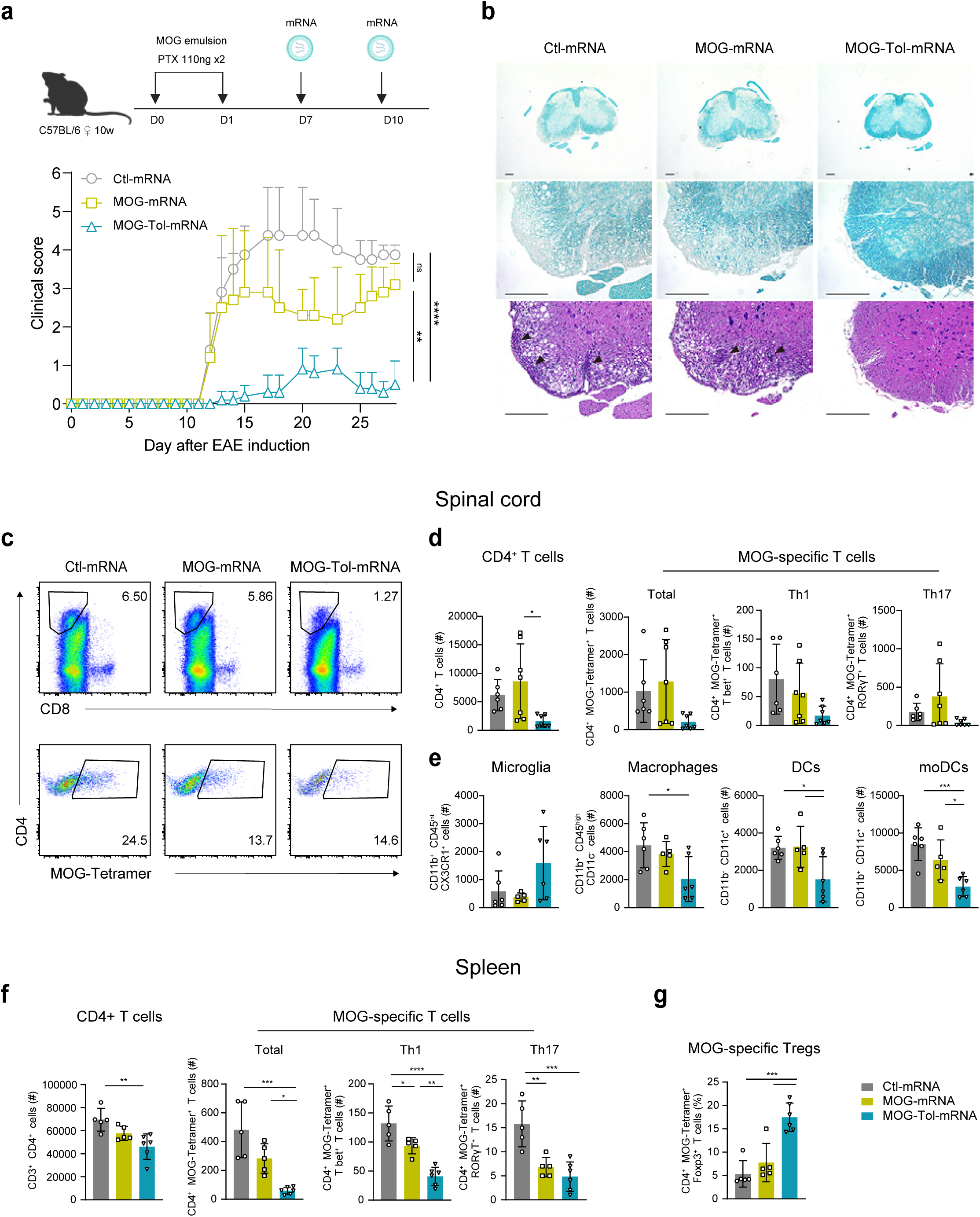
MOG-Tol-mRNA prevents disease progression in EAE. (a) Clinical disease scores of EAE mice treated with mRNA at days 7 and 10 post-induction were monitored until day 28. (b) Representative images of spinal cord sections at day 17 post-EAE induction. Inflammatory cell infiltration was assessed by hematoxylin and eosin (H&E) staining, and demyelination was evaluated by Luxol Fast Blue (LFB) staining. (c) The frequencies of CD4⁺ T cells and MOG–specific T cells infiltrating the spinal cord were analyzed by flow cytometry at day 28 post-EAE induction. (d) The numbers of total CD4⁺ T cells, as well as MOG–specific T-bet⁺ Th1 and RORγt⁺ Th17 cells, in the spinal cord were quantified at day 28 post-EAE induction. (e) The numbers of microglia, macrophages, dendritic cells (DCs), and monocyte-derived dendritic cells (moDCs) in the spinal cord were quantified at day 28 post-EAE induction. (f) The numbers of total CD4⁺ T cells, as well as MOG-specific T-bet⁺ Th1 and RORγt⁺ Th17 cells, in the spleen were analyzed at day 28 post-EAE induction. (g) The proportion of MOG-specific Foxp3⁺ Tregs in the spleen at day 28 post-EAE induction.

Interestingly, despite a reduction in MOG-specific T cells in the spleen, the proportion of antigen-specific Tregs was increased in the MOG-Tol-mRNA group (Fig. 2g). Evaluation of ICOS, PD-L1, CTLA-4, TIGIT, LAG3, and Tim-3 expression on antigen-specific Tregs revealed slight increases in PD-L1, TIGIT, and Tim-3 (Extended Data Fig. 5).

Remarkably, the MOG-Tol-mRNA conferred comparable prophylactic efficacy to existing MOG-mRNA vaccines^13^, despite containing only approximately one-third the antigen dose (Extended Data Fig. 6a). Compared with the 21 μg MOG-mRNA, the numbers of infiltrating CD4⁺ T cells and MOG-specific T cells were similar; however, inflammatory and Th17-associated cytokines were more effectively suppressed by the MOG-Tol-mRNA treatment (Extended Data Fig. 6b, c).

We next assessed the therapeutic efficacy of the mRNA treatment in mice that had already developed EAE symptoms. While MOG-mRNA alone significantly alleviated disease, co-administration with TGF-β and PD-L1 further enhanced the therapeutic effect. The therapeutic efficacy was also comparable to that of conventional 21 μg MOG-mRNA (Extended Data Fig. 7).

These findings suggest that treatment with the Tol-mRNA induces antigen-specific Tregs in the spleen, suppresses activation of MOG-specific T cells and other T cell subsets, and thereby prevents disease exacerbation in autoimmune pathology. The combination therapy using TGF-β and PD-L1 may offer a safer approach for treating autoimmune diseases.

### OVA-Tol-mRNA inhibits the development of allergic symptoms

We next investigated whether allergic responses to the food antigen OVA could be suppressed by mRNA treatment. BALB/c mice were pretreated by mRNA 14 days before the induction of OVA-induced food allergy (Fig. 3a). Notably, the OVA-Tol-mRNA prevented disease development in most treated mice. In contrast, mice treated with Ctl-mRNA or OVA-mRNA exhibited a similar level of hypothermia and severe diarrhea as untreated mice (Fig. 3b, c).

**Figure 3.**
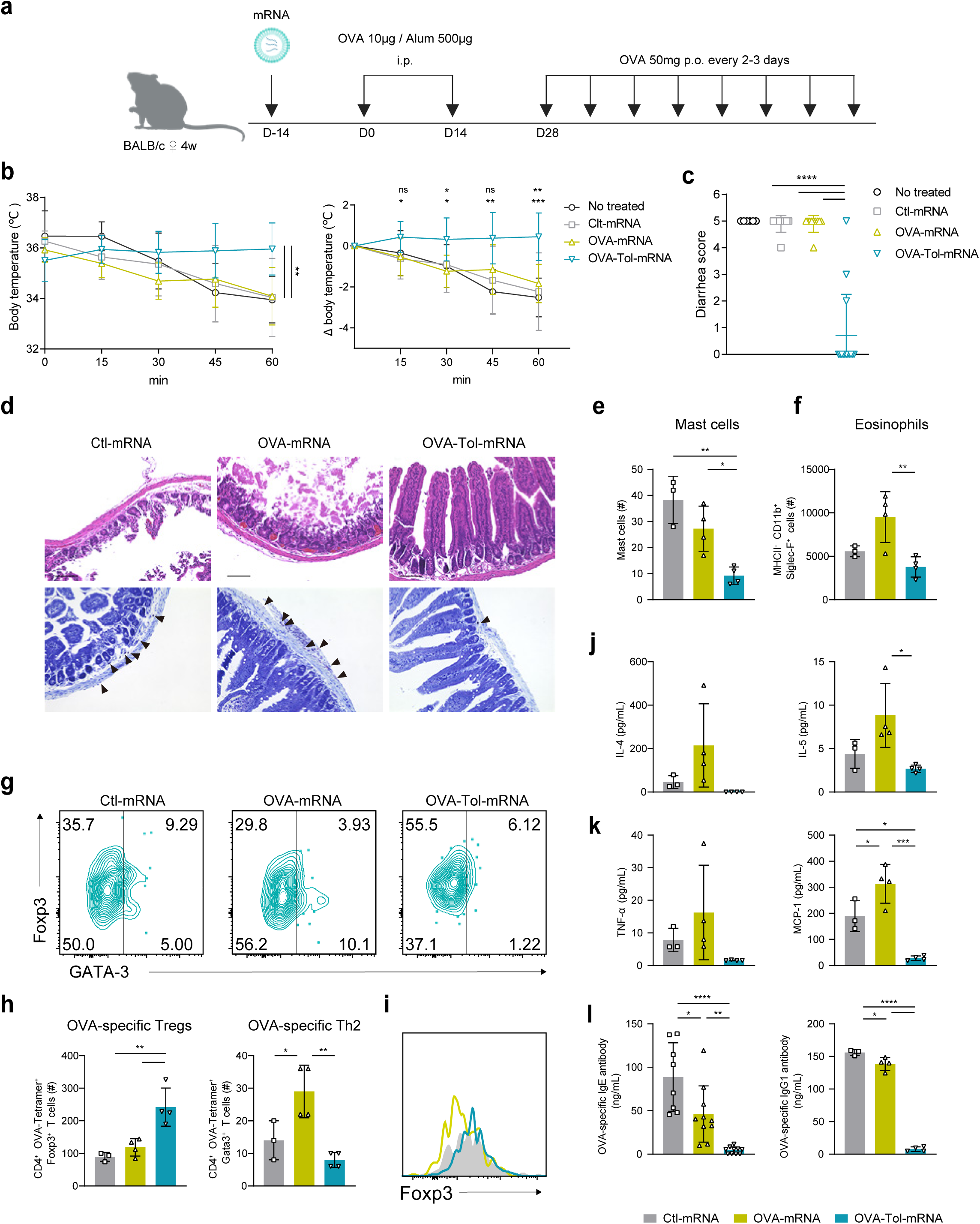
OVA-Tol-mRNA inhibits the development of allergic symptoms. (a) Schematic representation of the experimental design for induction of the OVA food allergy model and mRNA administration. (b) Rectal temperature was monitored for 60 min after the final administration of OVA protein, and the change in temperature over time and the maximum decrease in temperature were evaluated. (c) Diarrhea severity was evaluated after the final administration of OVA protein solution. (d) Representative images of jejunal sections at the end of the experiment. Jejunal architecture was assessed by H&E staining, and mast cell infiltration was evaluated by toluidine blue staining. (e) The number of mast cells in toluidine blue–stained jejunal sections (d) was quantified. (f) The number of eosinophils in the spleens of allergy-induced mice was quantified by flow cytometry. (g and h) At the study endpoint, the frequency and absolute number of OVA-specific Foxp3⁺ Tregs and GATA3⁺ Th2 cells in the spleens of allergy-induced mice. (i and j) Serum levels of Th2-associated and inflammatory cytokines, including IL-4, IL-5, TNF-α, and MCP-1, were determined at the end of the experiment. (k) In parallel, OVA-specific humoral responses were evaluated by measuring IgE and IgG1 antibody titers in the serum.

**Figure 4.**
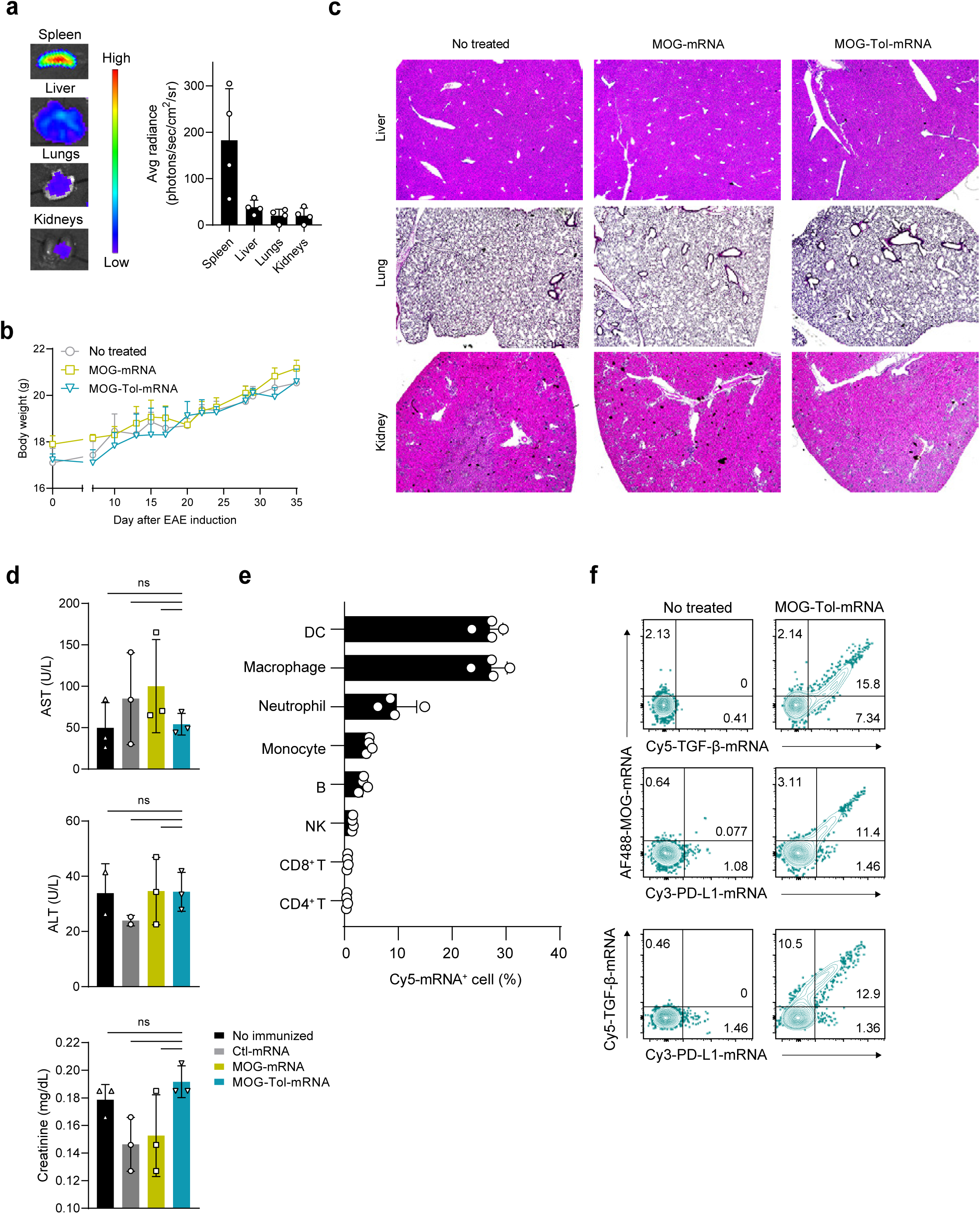
Cellular uptake and co-delivery of multiple mRNAs in the spleen with preserved systemic safety. (a) The biodistribution of administered mRNA was evaluated *in vivo* using Akaluc-Venus–encoding mRNA. (b) Body weight was measured for 35 days after mRNA administration. (c) Representative H&E–stained images of the liver, lung, and kidney collected after mRNA administration. Tissue morphology was assessed to evaluate potential histopathological changes associated with treatment. (d) Serum levels of aspartate aminotransferase (AST), alanine aminotransferase (ALT), and creatinine were measured to assess liver and kidney function. (e) Identification of mRNA-uptaking cell populations in the spleen using Cy5-labeled mRNA. (f) Co-localization of Alexa Fluor 488–labeled MOG-mRNA, Cy5-labeled TGF-β-mRNA, and Cy3-labeled PD-L1-mRNA in splenocytes was evaluated by flow cytometry.

Histological analysis revealed that the villus architecture in the jejunum was preserved in the OVA-Tol-mRNA-treated group (Fig. 3d). Infiltration of mast cells into the lamina propria was significantly suppressed in the OVA-Tol-mRNA-treated mice (Fig. 3d, e). As eosinophil accumulation is a hallmark of allergic patients^18^, we also examined eosinophil infiltration. The OVA-Tol-mRNA significantly reduced eosinophil accumulation (Fig. 3f).

Interestingly, while OVA-specific Th2 cell frequencies and numbers increased in mice treated with Ctl-mRNA or OVA-mRNA, no such increase was observed in the OVA-Tol-mRNA. Additionally, treatment with the OVA-Tol-mRNA led to increased antigen-specific Tregs (Fig. 3g, h).

Serum levels of Th2-associated cytokines were also significantly reduced in the OVA-Tol-mRNA group, consistent with the reduction in OVA-specific Th2 cells (Fig.3i). Moreover, inflammatory cytokines such as TNF-α and MCP-1 were significantly reduced (Fig.3j). In food allergy, production of antigen-specific IgE and IgG is closely associated with disease severity^19^. The OVA-Tol-mRNA significantly suppressed both OVA-specific IgE and IgG1 production (Fig.3k).

These findings suggest that antigen-specific Tregs may play a central role in suppressing allergic responses.

### Both PD-L1 and TGF-β are required for suppression of allergic responses

To dissect the contributions of PD-L1 and TGF-β in allergy suppression, we evaluated the effects of combining OVA-mRNA with either PD-L1 mRNA or TGF-β mRNA. The combination of OVA-mRNA and TGF-β-mRNA failed to prevent disease development. In contrast, the OVA-mRNA and PD-L1-mRNA exhibited partial protection, but full protection was not achieved (Extended Data Fig. 8a, b). Interestingly, serum levels of OVA-specific IgE were suppressed in the TGF-β combination group (Extended Data Fig. 8c). Expansion of OVA-specific Tregs was only observed when both PD-L1 and TGF-β were included. Likewise, reduction of OVA-specific Th2 cells was only achieved in the OVA-Tol-mRNA (Extended Data Fig. 8d). Consistent with this, Th2- and inflammation-associated cytokine production was higher in groups receiving only single immunoregulatory molecules compared to the OVA-Tol-mRNA group (Extended Data Fig. 8e).

Taken together, these results demonstrate that PD-L1 and TGF-β, acting through distinct yet complementary pathways, are essential for antigen-specific Treg induction and effective suppression of allergic inflammation, underscoring the importance of their combination with antigen mRNA for preventing allergic disease.

### Tol-mRNA mediates targeted delivery to splenic APCs while preserving antigen-specific immune responses and safety

To evaluate the *in vivo* biodistribution of mRNA, mice were intravenously administered Akaluc-Venus–encoding mRNA formulated with LPX at an RNA:LPX charge ratio of 2:1.7. Consistent with previous reports, the majority of the administered mRNA was preferentially delivered to the spleen^15^, with lower but detectable signals observed in the lungs, kidneys, and liver (Fig.4a). To evaluate the safety of systemic mRNA administration, C57BL/6 mice were treated twice with mRNA. mRNA-treated mice exhibited normal body weight gain comparable to untreated controls (Fig.4b). Because biodistribution studies detected signals not only in the spleen but also in the lungs, kidneys, and liver, these organs were examined for toxicity. Histological analysis revealed no apparent tissue injury or inflammatory cell accumulation (Fig.4c), and serum AST, ALT, and creatinine levels were not significantly altered compared with controls, indicating no overt hepatic or renal toxicity under the conditions tested (Fig.4d). To further identify the splenic cell populations responsible for mRNA uptake, mice were intravenously injected with Cy5-labeled mRNA. Flow cytometric analysis demonstrated that macrophages and DCs constituted the primary mRNA-uptaking populations (Fig.4e). Importantly, most Alexa Fluor 488-MOG-mRNA, Cy3-PD-L1-mRNA and Cy5-labeled TGF-β-mRNAs were co-localized within the same cells, indicating that multiple distinct mRNAs can be simultaneously delivered to individual cells (Fig.4f). Collectively, these results support the potential of the Tol-mRNA platform to recapitulate tolerogenic DC–like signaling *in vivo* through coordinated delivery of antigenic and immunoregulatory mRNAs to individual antigen-presenting cells.

To test for generalized immune suppression, mice in which MOG-specific tolerance had been established were subsequently immunized with OVA. Across all prior mRNA treatment conditions, OVA immunization induced comparable expansion of OVA-specific T cells, and the expanded cells displayed an effector phenotype with cytotoxic mediator expression (Extended Data Fig. 9a, b). *In vivo* cytotoxicity assays further confirmed preserved functional killing, as OVA peptide-loaded target cells were selectively eliminated (Extended Data Fig. 9c). These results indicate that systemic mRNA treatment is well tolerated and does not cause nonspecific immune activation, supporting the safety and antigen specificity of this approach.

### The humanized Tol-mRNA selectively suppresses antigen-specific T cells and induces Tregs

We next sought to determine whether the human Tol-mRNA suppresses antigen-specific T cell responses and promotes the induction of antigen-specific Tregs in a human experimental setting. HLA-DRB1*15:01 is one of the HLA class II alleles most strongly associated with human multiple sclerosis ^20,21^. In this study, we employed TCR-Ob.2F3, a myelin-reactive T cell receptor restricted to MBP₈₅_–_₉₉–HLA-DRB1*15:01^22,23^.

293T cells lacking β2m were transfected with mRNAs encoding MBP₈₅_–_₉₉–HLA-DRB1*15:01 alone or together with human TGF-β and human PD-L1. These cells were then co-cultured with TCR-Ob.2F3–expressing T cells (Fig.5a). While MBP–HLA alone induced a dose-dependent expansion of TCR-Ob.2F3^+^ T cells, MBP-Tol-mRNA markedly suppressed this expansion (Fig.5b, c). Consistently, inflammatory cytokines in the culture supernatants were significantly reduced (Fig.5d). Moreover, MBP-Tol-mRNA selectively induced differentiation of TCR-Ob.2F3^+^ T cells into Tregs, whereas TCR-Ob.2F3^-^ T cells were largely unaffected (Fig.5e).

We next examined whether the MBP-Tol-mRNA could suppress already activated T cells. To this end, TCR-T cells were pre-activated with CD3/CD28 beads 24 hours prior to co-culture (Extended Data Fig. 10a). Even under these conditions, MBP-Tol-mRNA strongly suppressed the activation of TCR-T cells. Notably, this suppressive effect was restricted to the TCR^+^ population, while TCR^-^ T cells exhibited activation levels comparable to control conditions (Extended Data Fig. 10b).

Together, these results demonstrate that the human Tol-mRNA delivers immunosuppressive signals in an antigen-specific manner, enabling selective immune regulation and preferential induction of antigen-specific Tregs.

### The humanized Tol-mRNA suppresses antigen-specific CD8⁺ T cells

Autoreactive CD8⁺ T cells have been reported to infiltrate target tissues and directly mediate tissue damage in many autoimmune diseases^24^. To determine whether the Tol-mRNA can suppress antigen-specific CD8⁺ T cell responses, we generated NY-ESO-1–specific T cells from HLA-A*02:01–positive donors and co-cultured with mRNA transfected 293T cells (Fig. 5f). NY-ESO-1–HLA-expressing 293T cells strongly expanded NY-ESO-1-specific T cells. In contrast, the NY-ESO-1-Tol-mRNA significantly suppressed the proliferation of NY-ESO-1-specific T cells and reduced expression of CD69 (Fig. 5g, Extended Data Fig. 11a). This effect persisted even when T cells were pre-activated (Fig. 5h). Importantly, NY-ESO-1-negative T cells showed comparable proliferation under all conditions, suggesting antigen specificity for this inhibitory effect (Extended Data Fig. 11b).

**Figure 5.**
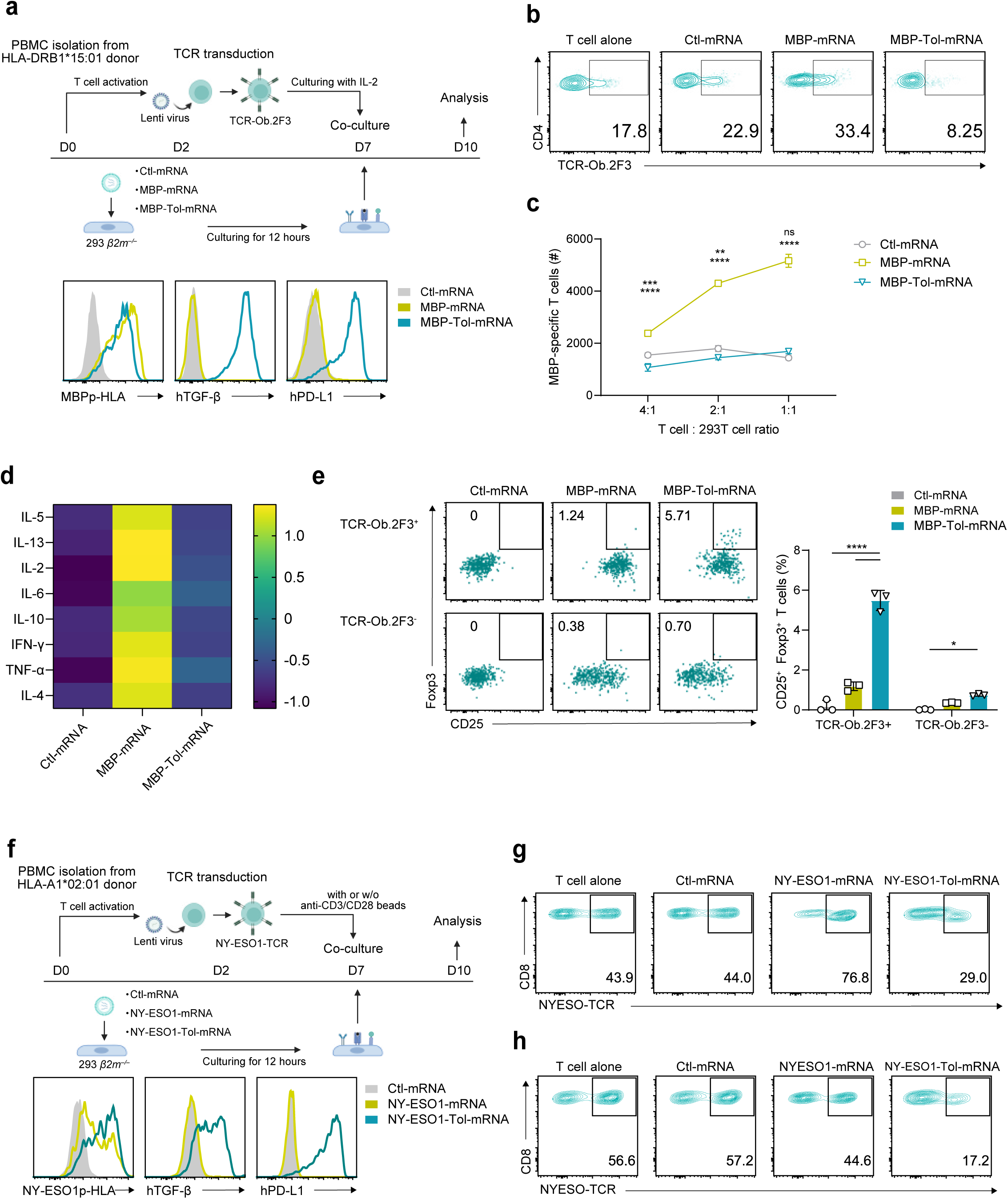
The humanized Tol-mRNA selectively suppresses antigen-specific T cells and induces Tregs. (a) Schematic representation of the experimental protocol using human TCR-T cells, together with expression of MBP-HLA, TGF-β, and PD-L1 in 293T cells. (b) The frequency of TCR-Ob.2F3–expressing cells within human CD4⁺ T cells was analyzed by flow cytometry after 72 h of co-culture. (c) The number of MBP-specific T cells was quantified after co-culture with mRNA-transfected 293T cells at different T cell–to–293T cell ratios (4:1, 2:1, and 1:1). (d) Cytokine levels in the culture supernatants were analyzed and visualized as a Z-score–normalized histogram. Relative changes in cytokine production across conditions were calculated by standardizing each cytokine to its mean and standard deviation, allowing comparison of expression patterns independent of absolute concentration differences. (e) At 96 h post-culture, the frequencies of Foxp3⁺ and CD25⁺ Tregs were analyzed within TCR-Ob.2F3⁺ and TCR-Ob.2F3⁻ CD4⁺ T cell populations. (f) Schematic representation of the experimental protocol using human CD8⁺ TCR-T cells, together with expression of NY-ESO-HLA, TGF-β, and PD-L1 in 293T cells. (g) The frequency of NY-ESO–specific TCR-expressing cells within human CD8⁺ T cells was analyzed after 72 h of co-culture. (h) TCR-T cells were stimulated with CD3/CD28 beads for 24 h prior to co-culture with 293T cells, followed by analysis as described in (g).

Thus, the humanized Tol-mRNA provides a strategy to selectively suppress pathogenic antigen-specific CD8⁺ T cells, supporting its potential utility for antigen-specific immune modulation in autoimmune diseases.

## Discussion

In this study, we show that robust antigen-specific immune tolerance can be induced *in vivo* by delivering immunoregulatory signals together with antigen presentation. We developed a three-component construct, Tol-mRNA, which integrates an antigen with PD-L1 and TGF-β, and found that it efficiently induces antigen-specific Tregs across multiple antigen contexts while suppressing inflammatory T cell responses. A key advance is that the mRNA platform enables reconstruction of the integrated signals required for physiological peripheral Treg induction, which is difficult to achieve with antigen alone^11,12^.

Tol-mRNA consistently induced tolerance in distinct disease settings, including autoimmunity and allergy. Importantly, previous studies have shown that antigen alone can confer protection in several mouse models of multiple sclerosis ^13^, and our observations are consistent with the notion that increasing antigen exposure can bias responses toward reduced pathogenicity. However, Tol-mRNA provides several conceptual and practical advantages over antigen-only strategies. First, Tol-mRNA achieves robust tolerance with a substantially lower effective antigen dose, suggesting a wider therapeutic window and potentially improved manufacturability and safety. Second, because peripheral Treg induction physiologically requires antigen recognition in the context of inhibitory and differentiation cues, the coordinated delivery of PD-L1 and TGF-β together with antigen provides a principled mechanism to promote Foxp3-associated programs and maintain suppressive function, rather than relying on dose-dependent effects of antigen presentation alone. Third, by enforcing co-expression of antigen and immunoregulatory signals within the same antigen-presenting cell, Tol-mRNA is designed to selectively deliver suppressive cues to cognate T cells at the time of antigen recognition, thereby minimizing bystander immunosuppression while more efficiently redirecting pathogenic immunity toward tolerance.

The humanized Tol-mRNA experiments further support the potential translational relevance of this platform by demonstrating antigen-dependent suppression of human CD4⁺ and CD8⁺ T-cell responses and preferential induction of antigen-specific Tregs. Because the human experiments used engineered antigen-presenting cells and TCR-transduced T cells, future studies using primary human APCs and patient-derived autoreactive T cells will be important to validate translational relevance.

mRNA therapeutics are already clinically established and effective in infectious disease and cancer settings^25–27^. For autoimmune applications, both immunological precision and systemic safety are critical. Recent low-immunogenic mRNA platforms based on improved synthesis chemistry and sequence engineering could mitigate the transient immune activation observed with our current formulation and thereby widen the safety margin^28,29^. In parallel, emerging low-inflammatory LNP formulations may enable safer delivery while preserving efficient targeting^30–33^. Finally, extracellular vesicles represent a promising alternative delivery modality with potentially lower immunogenicity and favorable biocompatibility ^34,35^.

In summary, Tol-mRNA integrates antigen presentation with PD-L1 and TGF-β signaling to induce antigen-specific immune tolerance *in vivo* and suppress autoimmune and allergic disease phenotypes. The ability to reconstruct multiple tolerance-inducing signals using a scalable mRNA modality provides a foundation for antigen-specific immunotherapies aimed at durable disease modification.

## Materials and Methods

### Cell lines

Human embryonic kidney (HEK)-293T cells (ATCC, Cat# CRL-3216) and Lenti-X cells (Takara, 632180) were cultured in Dulbecco’s modified Eagle’s medium (DMEM; Thermo Fisher Scientific, 08458-16) supplemented with 10% heat-inactivated fetal calf serum (FCS; Thermo Fisher Scientific, A5256701), 100 U/mL penicillin, and 100 U/mL streptomycin (FUJIFILM Wako, 16823191). Murine lymphocytes were cultured in RPMI (Nacalai Tesque, 30264-56) supplemented with 10% FCS, 1× non-essential amino acid (FUJIFILM Wako, 139-15651), 1 nmol/L sodium pyruvate (Nacalai Tesque), 100 U/mL penicillin, 100 U/mL streptomycin, and 0.05 μM 2-mercaptoethanol (Thermo Fisher Scientific, 21985023). Human PBMCs were resuspended in X-VIVO 15 medium (Lonza, 04-418Q) supplemented with 5% human AB serum (FUJIFILM Wako, 553-21741) and stimulated in 24-well plates with human CD3/CD28 beads (Gibco, 11131D) and recombinant human IL-2 (Biolegend, 589104) at 20 ng/mL.

### Mice

C57BL/6 and BALB/c mice were purchased from Japan SLC (Shizuoka, Japan) and The Jackson Laboratory (Bar Harbor, ME, USA). 2D2 TCR transgenic mice (expressing a T cell receptor specific for the MOG_35–55_ peptide presented by I-Ab)^36^ were maintained on a C57BL/6 background and bred in our facility.

All mice were maintained under specific pathogen-free facility (SPF) conditions at the animal facility of Kanazawa university.

### RNA construction

*In vitro* transcription of RNA constructs was performed using a plasmid backbone containing a T7 promoter, 5′ untranslated region (5′UTR), 3′ untranslated region (3′UTR), and a poly(A) tail of 129 nucleotides.

As antigens, we used the following peptide sequences: amino acids 315–347 of OVA, amino acids 27–63 of MOG, a hybrid sequence referred to as HIP2.5 (consisting of residues 43–56 of mouse proinsulin-2 and residues 358–370 of chromogranin A [WE14])^37^, amino acids 1–31 of insulin β-chain. To enhance antigen processing and presentation, each antigen sequence was flanked by sequences encoding an MHC class I signal peptide at the N-terminus and an MHC class I trafficking domain (MITD) at the C-terminus^38^.

For control immunization with mRNA, constructs encoding either an empty MITD (lacking any antigen sequence) or a fusion protein of Venus and Akaluc were used^39^. Additional constructs encoding mouse immunomodulatory molecules—TGF-β-CD8 (C33S, C223S, C225S)^40–42^, PD-L1, CD80, OVA whole sequence, HLA-DR2 alpha chain (F12S, M23K)^23^, MBP-HLA-DRB1*15:01 (P11S)^23^, active hTGF-β-CD8 (C33S, C223S, C225S)^40–42^, hPD-L1 and NY-ESO-1–HLA-A*02:01—were also designed using the same plasmid backbone. All amino acid sequences are shown in Extended Data Table 1.

### mRNA synthesis

All mRNAs were synthesized by *in vitro* transcription using the HiScribe™ T7 mRNA Kit (New England Biolabs, E2080S), incorporating N1-methyl-pseudouridine (TriLink Bio, N-1081-10) in place of uridine^43,44^.

Transcribed RNAs were purified and eluted in nuclease-free H_2_O, and stored at –80□°C until use. The integrity and purity of synthesized mRNA were confirmed by analysis using an Agilent TapeStation system (Agilent Technology).

### Preparation of lipoplex (LPX) lipid solution

A lipid solution was prepared by mixing 1,2-di-O-octadecenyl-3-trimethylammonium propane (DOTMA; Avanti, 890898P) and 1,2-dioleoyl-sn-glycero-3-phosphoethanolamine (DOPE; Avanti, 850725P-25MG) at a molar ratio of 2:1, resulting in a final concentration of 4.66 mM DOTMA and 2.33 mM DOPE in water. The lipid mixture was processed using an extruder to produce small unilamellar vesicles, which were used as LPX liposomes^45^.

RNA–lipoplexes (RNA–LPX) were prepared under sterile and RNase-free conditions. The final solution contained 150 mM NaCl. The molar ratio of RNA to DOTMA was adjusted to 2.0:1.7^15^.

### Preparation of lipid nanoparticle (LNP)

LNPs encapsulating mRNA were formulated using a mixture of lipid components consisting of 0.5–15% polyethylene glycol (PEG)-modified lipid, 5–25% non-cationic lipid, 25–55% sterol, and 20–60% ionizable cationic lipid. The PEG-modified lipid used was 1,2-dimyristoyl-rac-glycero-3-methoxypolyethylene glycol-2000 (DMG-PEG 2000; Avanti, 880151P), the non-cationic lipid was 1,2-distearoyl-sn-glycero-3-phosphocholine (DSPC; Avanti, 850365P), the sterol was cholesterol, and the ionizable cationic lipid was heptadecan-9-yl 8-[(2-hydroxyethyl)(6-oxo-6-(undecyloxy)hexyl)amino]octanoate (SM-102; NARD Institute, Ltd., Kobe, Japan) ^46^.

### Antibodies

Antibody staining was performed according to standard protocols. Monoclonal antibodies used for surface staining included the following: anti-mCD3 (17A2), mCD4 (GK1.5), mCD8 (53–6.7), mCD11b (M1/70), mCD11c (N418), mCD19 (6D5), mCD25 (PC61), mPD-L1 (10F.9G2), mLAG-3 (C9B7W), mTim-3 (B8.2C12), mTIGIT (1G9), mNeuropilin-1 (3E12), mICOS (7E.17G9), mCCR6 (29-2L17), mCXCR3 (CXCR3-173), mCD103 (W19396D), mTCR-Vα2 (B20.1), mTCR-Vβ5 (MR9-4), mTCR-Vα3.2 (RR3-16), mTCR-Vβ11 (KT11), mTCR β chain (H57-597), mCD44 (IM7), mCD45 (30-F11), mCD45.1 (A20), mCD45.2 (104), mCD62L (MEL-14), mCD69 (H1.2F3), mCD80 (16-10A1), CD86 (A17199A), mNK1.1 (S17016D), mLY6G (1A8), mF4/80 (BM8), mSiglec F (S17007L), mIL-2 (JES6-5H4), mTGF-β (TW7-16B4), HLA-DR2 (Tü36), hTGF-β (TW4-9E7; BD Biosciences), hPD-L1 (MIH3), hCD25 (M-A251), hCD69 (FN50) .

Intracellular staining was performed using anti-mFoxp3 (MF-14), mCTLA-4 (UC10-4B9), mHelios (22F6), mT bet (4B10), mGATA-3 (16E10A23), mRORγT (Q3-378; BD Biosciences), mIgE (RME-1), mGranzyme B (QA16A02), mIFN-γ(XMG1.2), hFoxp3 (206D) antibody, in combination with the True-Nuclear™ Transcription Factor Buffer Set (BioLegend). Unless stated otherwise, items were purchased from Biolegend.

### Preparation of single cell suspension

Cells were isolated from the spleen, lymph nodes, and spinal cord of mice. Spleens were harvested and immediately placed in phosphate-buffered saline (PBS; FUJIFILM Wako, 045-29795), then mechanically dissociated on microscope slides. Red blood cells were lysed by incubation with 1□mL of ACK lysis buffer (Thermo Fisher Scientific, A10492-01).

Lymph nodes were excised, transferred into PBS, and then mechanically disrupted using a syringe plunger on a 100□μm filter mounted on a 50□mL tube. The cell suspensions were centrifuged at 500 × *g* for 3 minutes, the supernatant was carefully removed, and the cells were resuspended in 10□mL of FACS buffer.

Spinal cords were dissected and enzymatically digested using the Multi Tissue Dissociation Kit (Miltenyi Biotec, 130-110-201) following the manufacturer’s protocol. Cellular debris was eliminated using the Debris Removal Solution (Miltenyi Biotec, 130-109-398), and red blood cells were lysed using the ACK lysis.

### *In vivo* induction of antigen-specific Treg

For the *in vivo* induction of antigen-specific Treg expansion, mRNA was mixed with Invivo-jet RNA+ (Polyplus, 101000122) according to the manufacturer’s instructions. The following mRNA formulations were used: Peptide-Tol-mRNA (7□μg of Peptide-mRNA, 7□μg of TGFβ-CD8 mRNA, and 7□μg of PD-L1 mRNA), Peptide-mRNA alone (7□μg of Peptide-mRNA), or irrelevant mRNA (21□μg of Venus-Akaluc mRNA).

For LPX or LNP formulations, the following were used: MOG-Tol-mRNA (7□μg of MOG-mRNA, 7□μg of TGFβ-CD8 mRNA, and 7□μg of PD-L1 mRNA), MOG-mRNA (7□μg of MOG-mRNA and 14□μg of irrelevant mRNA), or 21□μg of irrelevant mRNA. These mRNAs were administered intravenously, and spleens were harvested 7 days post-injection.

A total of 3 × 10□ isolated cells were incubated in complete RPMI medium containing 5□μg/mL peptide–MHC class II tetramers (provided by the NIH Tetramer Core Facility) at 37□°C for 1 hour. Following tetramer staining, the supernatant was discarded, and dead cell staining and surface staining were performed at 4□°C for 15 minutes. Subsequently, cells were fixed and subjected to intracellular staining. Flow cytometric analysis was performed using a CytoFLEX flow cytometer (Beckman Coulter), and data were analyzed with FlowJo software version 10.10.0.

### *In vitro* suppression assay

To directly assess the suppressive function of the induced antigen-specific Tregs, splenic CD4⁺ T cells were isolated from mice after mRNA administration using a CD4⁺ T Cell Isolation Kit (Miltenyi Biotec; 130-104-454). The enriched CD4⁺ T cells were then stained with an APC-labeled peptide–MHC class II tetramer, and antigen-specific T cells were further enriched using anti-APC MicroBeads (Miltenyi Biotec, 130-090-855). The purified antigen-specific T cells were used as suppressor cells and co-cultured with CellTrace™ Violet (CTV; Invitrogen, C34557)-labeled 2D2 responder T cells (CD45.1/2), along with wild-type splenocytes (CD45.1) as stimulator cells, in the presence of 15 μg/mL MOG_35−55_ peptide. Suppressor and responder cells were co-cultured at multiple suppressor-to-responder ratios. After 72 hours of culture, responder 2D2 T-cell proliferation was assessed by CTV dilution.

### EAE induction and mRNA treatment in C57BL/6 mice

EAE was induced in 9- to 11-week-old female C57BL/6 mice by subcutaneous injection of MOG_₃₅–₅₅_ peptide emulsion supplied in the Hooke Kits™ for EAE induction (Hooke Laboratories, EK-2110), according to the manufacturer’s instructions. On the day of immunization and again one day later, mice were intraperitoneally injected with 110□ng of pertussis toxin. Mice were weighed and clinically scored daily starting from day 10 post-immunization using the following scale: 0, no clinical signs; 1, reduced tail tone; 2, impaired righting reflex; 3, hind limb paresis; 4, complete hind limb paralysis; 5, hind limb paralysis with forelimb weakness; and 6, moribund or dead. Clinical scoring was performed in a blinded manner by investigators who were unaware of the treatment groups to reduce bias.

To evaluate the prophylactic immune effects, EAE mice received intravenous administration of LPX-formulated mRNAs on days 7 and 10 after immunization. The mRNA formulations included: (1) Tol-mRNA containing 7□μg of each mRNAs; (2) 7□μg of MOG-mRNA combined with 14□μg of irrelevant mRNA; (3) 21□μg of irrelevant mRNA; or (4) 21□μg of MOG-mRNA.

To assess the therapeutic efficacy, mice that had developed EAE symptoms (clinical score 1–2) were treated with the same LPX-formulated mRNA compositions.

### Induction of OVA-induced food allergy and prophylactic mRNA treatment in BALB/c mice

To induce a food allergy, 5- to 6-week-old female BALB/c mice were sensitized by intraperitoneal injection of 100□µg of OVA (grade V, Sigma-Aldrich, A5503-5G) emulsified with 1□mg of Alum adjuvant (G-Biosciences, 786-1215). The sensitization was performed twice at a 2-week interval. Two weeks after the second sensitization, mice were orally challenged every other day for a total of 8 times with 50□mg of crude OVA (Nakarai Tesque, 01205-84), suspended in PBS. All mice were fasted overnight prior to each oral challenge.

Following the oral administration, rectal temperatures were measured at 0, 15, 30, 45, and 60 minutes post-challenge. Fecal consistency was assessed using a 6-point clinical score: 0, normal stool; 1, soft stool; 2, loose stool; 3, mild diarrhea; 4, severe diarrhea; and 5, liquid stool.

To evaluate the prophylactic immune effects, mice were intravenously administered mRNAs 10 to 14 days prior to the initiation of disease induction. The mRNA formulations included: (1) Tol-mRNA containing 7□μg of OVA-mRNA, 7□μg of TGFβ-CD8-mRNA, and 7□μg of PD-L1-mRNA; (2) 7□μg of OVA-mRNA combined with 14□μg of irrelevant mRNA; or (3) 21□μg of irrelevant mRNA.

At the end of the experiment, mice were anesthetized, and blood was collected via cardiac puncture. Following euthanasia, spleens and small intestines were harvested for downstream analyses.

### Histological analysis

At day 17 following EAE induction, mice were perfused and fixed with 4% paraformaldehyde. After fixation, the L4–L5 spinal cord segments were dissected and embedded in paraffin. Tissue sections were cut at a thickness of 3□μm and subjected to H&E staining and LFB staining. These histological stains were performed by Bio-Pathology Institute Co., Ltd. (Oita, Japan).

For the allergic mouse model, jejunal tissues were collected at the end of the experiment and fixed overnight in 4% paraformaldehyde. After paraffin embedding, sections were cut at 3□μm thickness and stained with H&E and toluidine blue. H&E staining was conducted by the Technical Support Section of Kanazawa University (Kanazawa, Japan), while Toluidine Blue staining was performed by Tokushima Institute for Molecular Pathology (Tokushima, Japan). All histological images were acquired using the BZ-X (KEYENCE).

### Measurement of serum anti-OVA IgE and IgG1 antibodies

Following the final oral challenge, blood was collected from the cervical vein under anesthesia on the last day of the experiment. Blood samples were left at room temperature for 1 hour to allow clotting, followed by centrifugation at 3000□rpm for 10 minutes to isolate serum.

Serum levels of anti-OVA IgE and IgG1 antibodies were measured by enzyme-linked immunosorbent assay (ELISA) using the Mouse Serum Anti-OVA IgE Antibody Assay Kit (Iwaki Chemical Co., Ltd., 3010) and the Anti-Ovalbumin IgG1 (mouse) ELISA Kit (Cayman Chemical, 500830), respectively, in accordance with the manufacturers’ protocols.

### Cytokine detection

At day 28 after EAE induction, spleens and spinal cords were harvested from mice. 2□×□10□ splenocytes or the entire population of isolated spinal cord cells were stimulated overnight with 15□μg/mL of MOG₃₅_–_₅₅ peptide. After stimulation, the supernatants were collected for cytokine analysis. In the food allergy model, serum was obtained from mice at the end of the experiment as previously described.

T cell-associated cytokines were quantified using the LEGENDplex™ Mouse Th Cytokine Panel (12-plex) with VbP V03 (BioLegend, 741044), according to the manufacturer’s instructions. Data were acquired using a CytoFLEX flow cytometer and analyzed with the LEGENDplex™ Data Analysis Software Suite. Additionally, serum levels of MCP-1 were measured by ELISA using the Mouse MCP-1 ELISA Kit (Proteintech, KE10006).

### Bioluminescence imaging

To evaluate the *in vivo* biodistribution of mRNA, BALB/c mice were intravenously administered 21 μg of Akaluc-Venus-encoding mRNA^39^. At 6 hours after injection, 100 µL of 30 mM AkaLumine-HCl (FUJIFILM Wako, 016-26704) solution was administered intraperitoneally, and bioluminescence was acquired 10 minutes later using an IVIS Imaging System (Xenogen). Regions of interest were quantified as average radiance, expressed as photons s^-1^ cm^-2^ sr^-1^ and displayed on a color scale.

### Evaluation of mRNA uptake

To identify cell populations that internalized mRNA *in vivo*, fluorescently labeled mRNAs were generated. Briefly, during *in vitro* transcription, Alexa Fluor 488-(Invitrogen, C11403), Cy3- (GLPBIO, GB20116) or Cy5-labeled UTP (GLPBIO, GB20007) was substituted for 10% of the total UTP, and mRNA was synthesized as described above. The labeled mRNAs were then complexed with LPX and intravenously administered to mice. 1 hour after mRNA administration, uptake of mRNA by splenic cell populations was analyzed by flow cytometry.

### Safety evaluation of mRNA administration

To assess long-term *in vivo* toxicity associated with mRNA administration, mice received two intravenous injections at a 1-week interval. Body weight was monitored every other day or every 2 days for approximately 1 month. At 35 days after the first administration, blood was collected to measure serum AST, ALT, and creatinine levels. Mice were then perfused and fixed, and the lungs, kidneys, and liver were harvested for histological evaluation by H&E staining.

Serum AST, ALT, and creatinine were quantified using LabAssay AST (FUJIFILM Wako, 299-97601), LabAssay ALT (FUJIFILM Wako, 293-97501), and LabAssay Creatinine kits (FUJIFILM Wako, 291-93901), respectively, according to the manufacturers’ instructions.

### Assessment of antigen-specific immune tolerance by *in vivo* cytotoxicity assay

To evaluate the induction of antigen-specific immune tolerance, LPX-formulated mRNAs were intravenously administered to CD45.1/2 congenic C57BL/6 mice. The mRNA formulations included: (1) MOG-Tol-mRNA; (2) MOG-mRNA alone; and (3) irrelevant mRNA.

7 days after the mRNA treatment, mice received an intravenous injection of 20□μg of OVA-mRNA, encoding the SIINFEKL CD8 epitope. Another 7 days later, a total of 2□×□10□ splenocytes were adoptively transferred into the mice. These cells consisted of a 1:1 mixture of CD45.2⁺ splenocytes pulsed with OVA peptide (SIINFEKL) and CD45.1⁺ splenocytes pulsed with an irrelevant control peptide (RPL18).

18 hours after cell transfer, splenocytes were isolated from recipient mice, and the expansion of OVA-specific CD8⁺ T cells as well as the *in vivo* cytotoxicity against peptide-pulsed target cells was analyzed using a flow cytometer.

### Human PBMC isolation

Peripheral blood was collected from healthy donors who were positive for either HLA-DRB1*15:01 or HLA-A*02:01. PBMCs were isolated by density-gradient centrifugation using Ficoll (Cytiva, 17144002) in Leucosep tubes (Greiner, 227290). The PBMC layer was collected and transferred to a 15 mL tube, and washed for 3 times. Finally, PBMCs were resuspended in X-VIVO 15 medium for downstream applications.

### Lentivirus production

One day prior to transfection, Lenti-X 293T cells were seeded into 6-well plates at 1.0 × 10^6^ cells/well. Lentiviral particles were produced by PEI-mediated co-transfection of the packaging plasmid pCMV (2.66 µg), the envelope plasmid pMD2.G (0.345 µg), and 3 µg of the pHR vector (addgene, Plasmid #79121) encoding either TCR-Ob.2F3 or an NY-ESO-1-specific TCR (Extended Data Table 1). At 12 hours post-transfection, the medium was replaced with 1.5 mL of fresh complete culture medium. Viral supernatants were collected at 24 and 48 hours post-transfection, passed through a 0.45 µm syringe filter, and used immediately or stored at 4°C until use.

### TCR transduction of primary human T cells

Purified PBMCs were stimulated in 24-well plates with human CD3/CD28 beads and recombinant human IL-2 at 20 ng/mL for 48 hours at 37°C.

For lentiviral transduction, non-treated 24-well plates were coated with RetroNectin (Takara, T100B) at 20 µg/mL in PBS by adding 350 µL per well and incubating overnight at 4°C.

Lentiviral supernatants were added to RetroNectin-coated wells at 2 mL per well. Plates were wrapped and centrifuged at 2000 × g for 3 hours at 32°C to load viral particles. Activated T cells were then added at 5 × 10^5^ cells/well in 1 mL of IL-2 supplemented medium, followed by centrifugation at 800 × g for 30 min at 32°C. On the following day, culture medium was replaced with fresh IL-2 supplemented medium. Thereafter, cells were maintained with half-medium changes and cultured at a density of 1 × 10□ cells/mL.

### Evaluation of the human Tol-mRNA *in vitro*

7 days after TCR transduction, the generated TCR-T cells were co-cultured with 293T cells. In experiments requiring pre-activation before co-culture, TCR-T cells were stimulated with human CD3/CD28 beads for 24 hours, and the beads were removed prior to co-culture. 293T cells (β2m-/-) were transfected 12 hours before co-culture with either ctl-mRNA, peptide-HLA-mRNAs, or peptide-HLA-mRNAs together with human TGF-β- and human PD-L1-mRNAs using the TransIT-mRNA Transfection Kit (Takara, MIR2225), according to the manufacturer’s instructions. Before co-culture, transfected 293T cells were treated with mitomycin C (Nacalai tesque, 20898-21) at 10 µg/mL for 2 hours at 37°C. A total of 5 × 10^4^ T cells was co-cultured with 293T cells at multiple T cell to 293T ratios. Proliferation of TCR-positive and TCR-negative T cells and cytokine levels in culture supernatants were assessed at 72 hours using LEGENDplex™ HU Th1/Th2 Panel (8-plex) w/ VbP V02 (Biolegend, 741030). Antigen-specific Treg induction was evaluated at 96 hours.

### Statistical analysis

One-way analysis of variance (ANOVA) was used to compare differences among multiple groups. For comparisons between two groups, unpaired two-tailed Student’s t-tests were performed. All statistical analyses were performed using GraphPad Prism version 8.0 (GraphPad Software, San Diego, CA, USA). A p-value of less than 0.05 was considered statistically significant.

## Acknowledgements

We thank the NIH Tetramer Core Facility (NIH Contract 75N93020D00005 and RRID:SCR_026557) for providing MOG-MHC class II tetramer, OVA-MHC class II tetramer, insulin β-MHC class II tetramer, HIP2.5-MHC class II tetramer and OVA-MHC class I tetramer. We would like to express our sincere gratitude to Professor Osamu Hori for technical assistance with the EAE experimental system, and to Assistant Professor Yuka Nagata and Professor Ryo Suzuki for technical assistance with the allergy experimental system.

## Data Availability

Upon reasonable request, data can be obtained from the corresponding author.

## Author Contributions

Shota Imai: Writing, Visualization, Validation, Methodology, Conceptualization, Funding acquisition, Investigation, Formal analysis, Data curation. Naoto Nishida: Investigation., Ryo Maeno: Investigation., Ayano Ikebuchi: Investigation., Hitoshi Sasatsuki: Investigation, Formal analysis, Data curation., Risa Saito: Investigation, Formal analysis, Data curation., Kazutaka Matoba: Investigation, Conceptualization, Visualization, Supervision., Sadahiro Iwabuchi: Investigation, Formal analysis, Data curation., Hisamichi Naito: Methodology., Tomohiro Iba: Investigation, Formal analysis, Data curation., Kanto Nagamori: Investigation., Toan Van Le: Investigation., Iriya Fujitsuka: Investigation., Makie Ueda: Investigation., Rikinari Hanayama: Conceptualization, Funding acquisition, Supervision., Tomoyoshi Yamano: Writing, Investigation, Conceptualization, Visualization, Funding acquisition, Supervision.

## Ethics Approval Statement

The use of mice in this study was based on their suitability as a widely accepted model for immunological investigations, enabling precise analysis of T-cell responses in a controlled setting. Female mice aged 6 to 8 weeks and weighing between 18 and 22 g were used at the time of experimentation. Mice were group-housed (up to five per cage) with ad libitum access to standard rodent chow and water, and maintained in a controlled environment with regulated temperature, humidity, and a 12-hour light/dark cycle. To support animal welfare and reduce stress, environmental enrichment such as nesting materials and shelters was provided. All procedures involving animals were carried out in compliance with the ARRIVE guidelines (https://arriveguidelines.org/) to ensure ethical and humane treatment. At the conclusion of the experiments, mice were euthanized using CO₂ inhalation with a gradual fill rate, in accordance with AVMA guidelines. The study protocol was reviewed and approved by the Animal Experimentation Committee of Kanazawa University (Kanazawa, Ishikawa, Japan) under approval number AP-204131.

---

**Extended Data Figure 1. Immunogenic mRNA combinations drive the induction of Foxp3**⁻ **antigen-specific T cells.**

The proportion of antigen-specific CD4⁺ T cells present within Foxp3⁺ Tregs or Foxp3⁻ conventional T cells in the spleens of mice administered immunogenic mRNA combinations.

**Extended Data Figure 2. Phenotypic analysis of MOG-specific Tregs.**

Expression of Helios, Nrp-1, CD25, CD69, ICOS, Tim-3, LAG-3, TIGIT, CTLA-4, PD-L1, CCR6, CXCR3, and CD103 was analyzed in mRNA-induced MOG-specific Foxp3⁺ Tregs by flow cytometry.

**Extended Data Figure 3. MOG-Tol-mRNA ameliorates disease severity in EAE.**

(a) The peak clinical score of EAE was evaluated in mice treated with mRNA. (b) Body weight of EAE-induced mice was monitored over time up to day 28 post-induction. (c) Demyelination in the spinal cord was quantified at day 17 post-EAE induction. Spinal cord sections were stained with Luxol Fast Blue, and the extent of demyelination was evaluated by imageJ. (d) The expression levels (mean fluorescence intensity, MFI) of MHC class II and CD86 were analyzed in splenic myeloid cell populations, including microglia, macrophages, DCs, and moDCs to assess their activation status. (e) The numbers of total T cells, CD4⁺ T cells, CD8⁺ T cells, NK cells, NKT cells, neutrophils, macrophages, and DCs among 5 × 10□ splenocytes from EAE-induced mice.

**Extended Data Figure 4. Cytokine production in EAE mice**

Spinal cord–derived cells and splenocytes from EAE-induced mice were stimulated *ex vivo* with MOG₃₅_–_₅₅ peptide, and cytokine production was measured.

**Extended Data Figure 5. Phenotypic analysis of MOG-specific Tregs in EAE mice.**

Expression of Foxp3, ICOS, PD-L1, CTLA-4, TIGIT, LAG-3, and Tim-3 was analyzed in MOG-specific Tregs in the spleens of EAE-induced mice.

**Extended Data Figure 6. Comparison of therapeutic efficacy between conventional MOG mRNA and Tol-mRNA in EAE.**

(a) Clinical scores of EAE mice treated with 20μg of MOG-mRNA or Tol-mRNA were monitored over time. (b) The proportions of total CD4⁺ T cells and MOG-specific CD4⁺ T cells infiltrating the spinal cord after treatment with conventional MOG mRNA or Tol-mRNA. (c) Spinal cord–derived cells and splenocytes from EAE-induced mice were stimulated *ex vivo* with MOG₃₅_–_₅₅ peptide, and cytokine production was measured.

**Extended Data Figure 7. Evaluation of the therapeutic function of Tol-mRNA in EAE mice.**

EAE clinical scores were monitored over time after administration of Ctl-mRNA, MOG-mRNA, Tol-mRNA, or 21μg of MOG-mRNA following disease onset to compare therapeutic efficacy among treatment groups.

**Extended Data Figure 8. Comparative analysis of allergy suppression induced by antigen combined with PD-L1 or TGF-β versus Tol-mRNA.**

(a) Following the final administration of the OVA protein solution, rectal temperature was measured over a 60-minute period to evaluate changes in body temperature over time and the maximum drop in body temperature. (b) The severity of diarrhea was assessed following the final administration of the OVA protein solution. (c) OVA-specific humoral immune responses were evaluated by measuring serum IgE antibody titers. (d) The number of OVA-specific Foxp3⁺ Treg cells and GATA3⁺ Th2 cells in the spleens of allergy-induced mice. (e) Serum concentrations of Th2-associated and inflammatory cytokines, including IL-4, IL-5, and MCP-1 were measured at the end of the experiment.

**Extended Data Figure 9. Tol-mRNA induces antigen-specific tolerance while preserving independent immune responses.**

(a) Schematic overview of experimental design for the evaluation of antigen-specific immune tolerance. The expansion of OVA-specific CD8^+^ T cells was analyzed after induction of MOG-specific immune tolerance to evaluate antigen specificity of the tolerogenic response. (b) Granzyme B and IFN-γ production in OVA-specific CD8^+^ T cells. (c) *In vivo* cytotoxic activity of OVA-specific CD8^+^ T cells was evaluated using a target cell killing assay. Splenocytes loaded with OVA peptide or control RPL-18 peptide were adoptively transferred into recipient mice, and target cell elimination was analyzed by flow cytometry 18 h after transfer.

**Extended Data Figure 10. Evaluation of the suppressive effect of Tol-mRNA in pre-activated human CD4**⁺ **T cells.**

(a) Schematic representation of the experimental protocol using human TCR-T cells, together with expression of MBP-HLA, TGF-β, and PD-L1 in 293T cells. TCR-T cells were pre-stimulated with CD3/CD28 beads for 24 h prior to co-culture with transfected 293T cells to evaluate antigen-specific responses. (b) The expansion of TCR-Ob.2F3⁺ and TCR-Ob.2F3⁻ CD4⁺ T cells was evaluated after co-culture. (c) The number of MBP-specific T cells was quantified after co-culture with mRNA-transfected 293T cells at different T cell–to–293T cell ratios (4:1, 2:1, and 1:1).

**Extended Data Figure 11. Expansion and activation of NY-ESO–specific CD8**⁺ **T cells following co-culture with mRNA-transfected 293T cells.**

(a) The expansion of NY-ESO–specific TCR-expressing CD8⁺ T cells and TCR^-^CD8⁺ T cells were evaluated after co-culture with mRNA-transfected 293T cells. (b) NY-ESO–specific TCR-expressing CD8⁺ T cells were pre-activated with CD3/CD28 beads prior to co-culture with mRNA-transfected 293T cells. Following co-culture, the expansion of TCR^+^ and TCR^-^ CD8⁺ T cells and CD69 expression were quantified by flow cytometry.

