## Extended data Table for "Modular mRNA platform for *in vivo* antigen-specific immune tolerance"

Extended data Table 1

Amino acid sequences of components referenced in Extended data Table 1.

| Element Name | Sequence |
|---|---|
| MOG-MITD | MRVTAPRTLILLLSGALALTETWAGSGGSGGGGSGGSPGKNATGMEV<br>GWYRSPFSRVVHLYRNGKDQDAEQAPGGSLGGGGSGIVGIVAGLAVL<br>AVVVIGAVVATVMCRRKSSGGKGGSYSQAASSDSAQGSDVSLTA |
| OVA-MITD | MRVTAPRTLILLLSGALALTETWAGSGGSGGGGSGGISSAESLKISQAV<br>HAAHAEINEAGREVVGSAEAGGSLGGGGSGIVGIVAGLAVLAVVVIGAV<br>VATVMCRRKSSGGKGGSYSQAASSDSAQGSDVSLTA |
| HIP2.5-MITD | MRVTAPRTLILLLSGALALTETWAGSGGSGGGGSGGELGGSPGDLQTL<br>ALWSRMDQLAKELTAGGSLGGGGSGIVGIVAGLAVLAVVVIGAVVATV<br>MCRRKSSGGKGGSYSQAASSDSAQGSDVSLTA |
| Insulin-β-MITD | MRVTAPRTLILLLSGALALTETWAGSGGSGGGGSGGVKQHLCGPHLV<br>ERLYLVCGEEGFFYTPKSRGGSLGGGGSGIVGIVAGLAVLAVVVIGAVV<br>ATVMCRRKSSGGKGGSYSQAASSDSAQGSDVSLTA |
| Empty MITD | MRVTAPRTLILLLSGALALTETWAGSGGSGGGGSGGGGSLGGGGSGIV<br>GIVAGLAVLAVVVIGAVVATVMCRRKSSGGKGGSYSQAASSDSAQGSD<br>VSLTA |
| Active TGF-β-CD8<br>(C33S, C223S, C225S) | MPPSGLRLLPLLLPLPWLLVLTPGRPAAGLSTSKTIDMELVKRKRIEAI<br>RGQILSKLRLASPPSQGEVPPGPLPEAVLALYNSTRDRVAGESADPEPE<br>PEADYYAKEVTRVLMVDRNNAIYEKTKDISHSIYMFFNTSDIREAVPEP<br>PLLSRAELRLQRLKSSVEQHVELYQKYSNNSWRYLGNRLLTPTDTPE<br>WLSFDVTGVVRQWLNQGDGIQGFRFSAHSSSDSKDNKLHVEINGISP<br>KRRGDLGTIHDMNRPFLLLMATPLERAQHLHSSRHRRALDTNYCFSS<br>TEKNCCVRQLYIDFRKDLGWKWIHEPKGYHANFCLGPCPYIWSLDTQ<br>YSKVLALYNQHNPGASASPCCVPQALEPLPIVYYVGRKPKVEQLSNMI<br>VRSCKCSGGGGSVISNSVMYFSSVVPVLQKVNSTTTKPVLRTPSPVHP<br>TGTSQPQRPEDCRPRGSVKGTGLDFACDIYIWAPLAGICVALLLSLIITLI<br>CYHRSRKRVCKCPRPLVRQEGKPRPSEKIV |
| PD-L1 | MRIFAGIIFTACCHLLRAFTITAPKDLYVVEYGSNVTMECRFPVERELDL<br>LALVVYWEKEDEQVIQFVAGEEDLKPQHSNFRGRASLPKDQLLKGNA<br>ALQITDVKLQDAGVYCCIISYGGADYKRITLKVNAPYRKINQRISVDPAT<br>SEHELICQAEGYPEAEVIWTNSDHQPVSGKRSVTTSRTEGMLLNVTSS<br>LRVNATANDVFYCTFWRSQPGQNHTAELIIPELPATHPPQNRTHWVLL<br>GSILLFLIVVSTVLLFLRKQVRMLDVEKCGVEDTSSKNRNDTQFEET |

| | |
|---|---|
| CD80 | MACNCQLMQDTPLLKFPCPRLILLFVLLIRLSQVSSDVDEQLSKSVKD<br>KVLLPCRYNSPHEDESEDRIYWQKHDKVVLSVIAGKLKVWPEYKNRTL<br>YDNTTYSLIILGLVLSDRGTYSCVVQKKERGTYEVKHLALVKLSIKADF<br>STPNITESGNPSADTKRITCFASGGFPKPRFSWLENGRELPGINTTISQ<br>DPESELYTISSQLDFNTTRNHTIKCLIKYGDAHVSEDFTWEKPPEDPPD<br>SKNTLVLFGAGFGAVITVVVIVVIIKCFCKHRSCFRRNEASRETNNSLTF<br>GPEEALAEQTVFL |
| Venus-Akaluc | MVSKGEELFTGVVPILVELDGDVNGHKFSVSGEGEGDATYGKLTLKLI<br>CTTGKLPVPWPTLVTTLGYGLQCFARYPDHMKQHDFFKSAMPEGYV<br>QERTIFFKDDGNYKTRAEVKFEGDTLVNRIELKGIDFKEDGNILGHKLE<br>YNYNSHNVYITADKQKNGIKANFKIRHNIEDGGVQLADHYQQNTPIGD<br>GPVLLPDNHYLSYQSALSKDPNEKRDHMVLLEFVTAAGITLGMDELY<br>KGSMEDAKNIKKGPAPFYPLEDGTAGEQLHKAMKRYALVPGAIAFTD<br>AHIQVDVTYAEYFEMSVRLAEAMRRYGLNTNHRIVVCSENSSQFFMPV<br>LGALFIGVAVAPANDIYNERELLNSMGISQPTVVFVSKKGLRKVLNVQK<br>KLPIIRKIIIMDSKTDYQGFQSMYTFVTSHLPPSFNEYDFVPESFDRDK<br>TIALIMNSSGSTGLPKGVALPHRTACVRFSHARDPIFGYQNIPDTAILSV<br>VPFHHGFGMFTTLGYLICGFRVVLMYRFEEELFLRSLQDYKIQSALLVP<br>TLFSCLAKSTLIDKYDLSSLREIASGGAPLSKEVGEAVAKRFRLPGIRQG<br>YGLTETTNAVMITPEGDRKPGSVGKVVPFFEAKVVDLVTGKTLGVNQR<br>GELCVRGPMIMSGYVNNPEATNALIDKDGWLHSGDIAYWDEDEHFFIV<br>DRLKSLIKYKGYQVAPAELEGILLQHPYIFDAGVAGLPDDDAGELPAAV<br>VVLEHGKTMTEKEIVDYVASQVTTAKKLRGGVVFVDEVPRGSTGKLDA<br>RKIREILTKAKKDGKIAV |
| OVA whole sequence | MGSIGAASMEFCFDVFKELKVHHANENIFYCPIAIMSALAMVYLGAKD<br>STRTQINKVVRFDKLPGFGDSIEAQCGTSVNVHSSLRDILNQITKPNDV<br>YSFSLASRLYAEERYPILPEYLQCVKELYRGGLEPINFQTAADQARELIN<br>SWVESQTNGIIRNVLQPSSVDSQTAMVLVNAIVFKGLWEKTFKDEDTQ<br>AMPFRVTEQESKPVQMMYQIGLFRVASMASEKMKILELPFASGTMSM<br>LVLLPDEVSGLEQLESIINFEKLTEWTSSNVMEERKIKVYLPRMKMEEK<br>YNLTSVLMAMGITDVFSSSANLSGISSAESLKISQAVHAAHAEINEAGR<br>EVVGSAEAGVDAASVSEEFRADHPFLFCIKHIATNAVLFFGRCVSP |
| HLA-DR2　alpha chain<br>(F12S, M23K) | MAISGVPVLGFFIIAVLMSAQESWAIKEEHVIIQAESYLNPDQSGEFKFD<br>FDGDEIFHVDMAKKETVWRLEEFGRFASFEAQGALANIAVDKANLEIM<br>TKRSNYTPITNVPPEVTVLTNSPVELREPNVLICFIDKFTPPVVNVTWL<br>RNGKPVTTGVSETVFLPREDHLFRKFHYLPFLPSTEDVYDCRVEHWG<br>LDEPLLKHWEFDAPSPLPETTENVVCALGLTVGLVGIIIGTIFIIKGLRKS<br>NAAERRGPL |

| | |
|---|---|
| MBP-HLA-DRB1*15:01<br>(P11S) | MVCLKLPGGSCMTALTVTLMVLSSPLALSENPVVHFFKNIVTPRGGG<br>GSGGGGSGGGGSGDTRPRFLWQSKRECHFFNGTERVRFLDRYFYNQ<br>EESVRFDSDVGEFRAVTELGRPDAEYWNSQKDILEQARAAVDTYCRH<br>NYGVVESFTVQRRVQPKVTVYPSKTQPLQHHNLLVCSVSGFYPGSIEV<br>RWFLNGQEEKAGMVSTGLIQNGDWTFQTLVMLETVPRSGEVYTCQV<br>EHPSVTSPLTVEWRARSESAQSKMLSGVGGFVLGLLFLGAGLFIYFRN<br>QKGHSGLQPTGFLS |
| Active hTGF-β-CD8<br>（C33S, C223S, C225S） | MPPSGLRLLPLLLPLPWLLVLTPGRPAAGLSTSKTIDMELVKRKRIEAI<br>RGQILSKLRLASPPSQGEVPPGPLPEAVLALYNSTRDRVAGESADPEPE<br>PEADYYAKEVTRVLMVDRNNAIYEKTKDISHSIYMFFNTSDIREAVPEP<br>PLLSRAELRLQRLKSSVEQHVELYQKYSNNSWRYLGNRLLTPTDTPE<br>WLSFDVTGVVRQWLNQGDGIQGFRFSAHSSSDSKDNKLHVEINGISP<br>KRRGDLGTIHDMNRPFLLLMATPLERAQHLHSSRHRRALDTNYCFSS<br>TEKNCCVRQLYIDFRKDLGWKWIHEPKGYHANFCLGPCPYIWSLDTQ<br>YSKVLALYNQHNPGASASPCCVPQALEPLPIVYYVGRKPKVEQLSNMI<br>VRSCKCSGGGGSTTTPAPRPPTPAPTIASQPLSLRPEACRPAAGGAVH<br>TRGLDFACDIYIWAPLAGTCGVLLLSLVITLYCNHRNRRRVCKCPRPVV<br>KSGDKPSLSARYV |
| hPD-L1 | MRIFAVFIFMTYWHLLNAFTVTVPKDLYVVEYGSNMTIECKFPVEKQL<br>DLAALIVYWEMEDKNIIQFVHGEEDLKVQHSSYRQRARLLKDQLSLGN<br>AALQITDVKLQDAGVYRCMISYGGADYKRITVKVNAPYNKINQRILVVD<br>PVTSEHELTCQAEGYPKAEVIWTSSDHQVLSGKTTTTNSKREEKLFNV<br>TSTLRINTTTNEIFYCTFRRLDPEENHTAELVIPELPLAHPPNERTHLVIL<br>GAILLCLGVALTFIFRLRKGRMMDVKKCGIQDTNSKKQSDTHLEET |
| NY-ESO-1–HLA-A*02:01 | MSRSVALAVLALLSLSGLEASLLMWITQCGGGGSGGGGSGGGGSIQRT<br>PKIQVYSRHPAENGKSNFLNCYVSGFHPSDIEVDLLKNGERIEKVEHSD<br>LSFSKDWSFYLLYYTEFTPTEKDEYACRVNHVTLSQPKIVKWDRDMG<br>GGGSGGGGSGGGGSGSHSMRYFFTSVSRPGRGEPRFIAVGYVDDTQF<br>VRFDSDAASQRMEPRAPWIEQEGPEYWDGETRKVKAHSQTLRVDLGT<br>LRGYYNQSEAGSHTVQRMYGCDVGSDWRFLRGYHQYAYDGKDYIAL<br>KEDLRSWTAADMAAQTTKHKWEAAHVAEQLRAYLEGTCVEWLRRYL<br>ENGKETLQRTDAPKTHMTHHAVSDHEATLRCWALSFYPAEITLTWQR<br>DGEDQTQDTELVETRPAGDGTFQKWAAVVVPSGQEQRYTCHVQHEG<br>LPKPLTLRWEPSSQPTIPIVGIIAGLVLLGAVITGAVVAAVMWRRKSSDR<br>KGGSYTQAASSDSAQGSDVSLTACKV |

| | |
|---|---|
| TCR Ob.2F3 | MLLLLLLLGPGSGLGAVVSQHPSWVISKSGTSVKIECRSLDFQATTMF<br>WYRQFPKQSLMLMATSNEGSKATYEQGVEKDKFLINHASLTLSTLTV<br>TSAHPEDSSFYICSARDLTSGSLNEQFFGPGTRLTVLEDLRNVTPPKVS<br>LFEPSKAEIANKQKATLVCLARGFFPDHVELSWWVNGKEVHSGVSTD<br>PQAYKESNYSYCLSSRLRVSATFWHNPRNHFRCQVQFHGLSEEDKW<br>PEGSPKPVTQNISAEAWGRADCGITSASYHQGVLSATILYEILLGKATLY<br>AVLVSGLVLMAMVKKKNSGSGATNFSLLKQAGDVEENPGPMETLLGV<br>SLVILWLQLARVNSQQGEEDPQALSIQEGENATMNCSYKTSINNLQWY<br>RQNSGRGLVHLILIRSNEREKHSGRLRVTLDTSKKSSSLLITASRAADT<br>ASYFCATDATSGTYKYIFGTGTRLKVLANIQNPEPAVYQLKDPRSQDS<br>TLCLFTDFDSQINVPKTMESGTFITDKTVLDMKAMDSKSNGAIAWSN<br>QTSFTCQDIFKETNATYPSSDVPCDATLTEKSFETDMNLNFQNLSVM<br>GLRILLLKVAGFNLLMTLRLWSS |
| NY-ESO-1-specific TCR | MSIGLLCCAALSLLWAGPVNAGVTQTPKFQVLKTGQSMTLQCAQDM<br>NHEYMSWYRQDPGMGLRLIHYSVAEGITDQGEVPNGYNVSRSTTEDF<br>PLRLLSAAPSQTSVYFCASSYVGAAGELFFGEGSRLTVLEDLRNVTPP<br>KVSLFEPSKAEIANKQKATLVCLARGFFPDHVELSWWVNGKEVHSGV<br>STDPQAYKESNYSYCLSSRLRVSATFWHNPRNHFRCQVQFHGLSEED<br>KWPEGSPKPVTQNISAEAWGRADCGITSASYHQGVLSATILYEILLGKA<br>TLYAVLVSGLVLMAMVKKKNSGSGATNFSLLKQAGDVEENPGPMETL<br>LGLLILWLQLQWVSSKQEVTQIPAALSVPEGENLVLNCSFTDSAIYNLQ<br>WFRQDPGKGLTSLLLIQSSQREQTSGRLNASLDKSSGRSTLYIAASQP<br>GDSATYLCAVRPLYGGSYIPTFGRGTSLIVHPDIQNPEPAVYQLKDPRS<br>QDSTLCLFTDFDSQINVPKTMESGTFITDKTVLDMKAMDSKSNGAIA<br>WSNQTSFTCQDIFKETNATYPSSDVPCDATLTEKSFETDMNLNFQNL<br>SVMGLRILLLKVAGFNLLMTLRLWSS |
