## Extended data Figures for "Modular mRNA platform for *in vivo* antigen-specific immune tolerance"

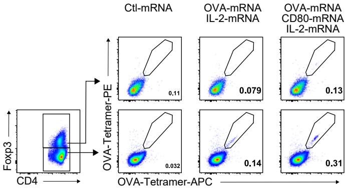


### Extended Data Figure 1

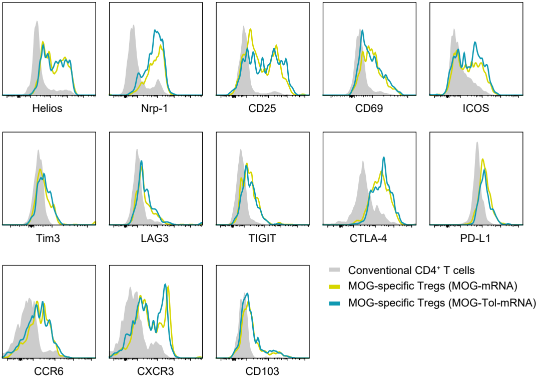


### Extended Data Figure 2

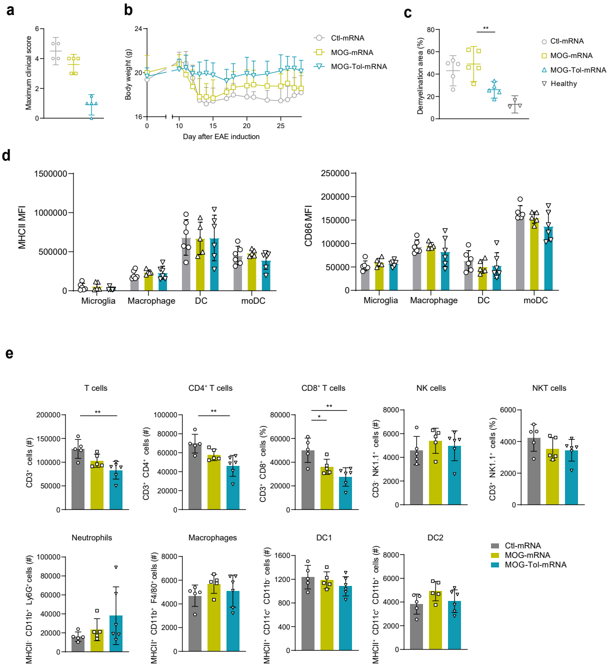


### Extended Data Figure 3

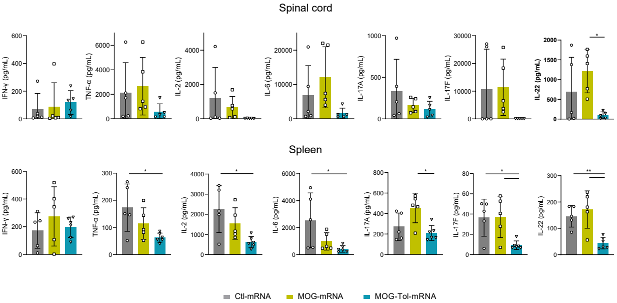


### Extended Data Figure 4

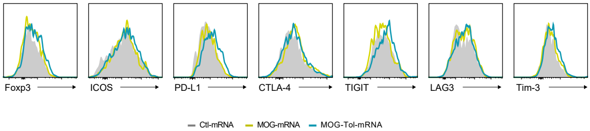


### Extended Data Figure 5

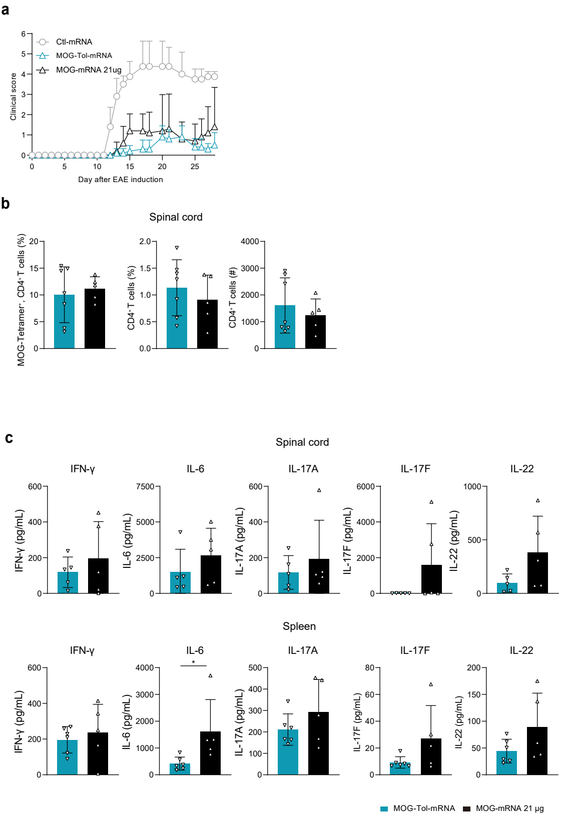


### Extended Data Figure 6

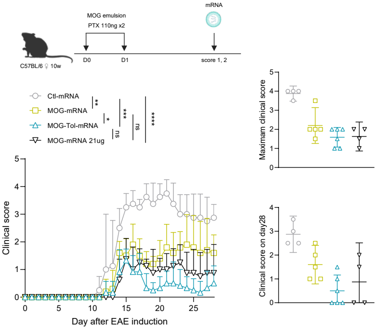


### Extended Data Figure 7

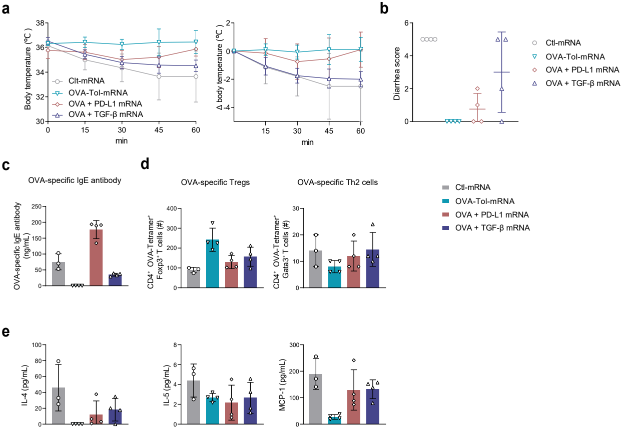


### Extended Data Figure 8

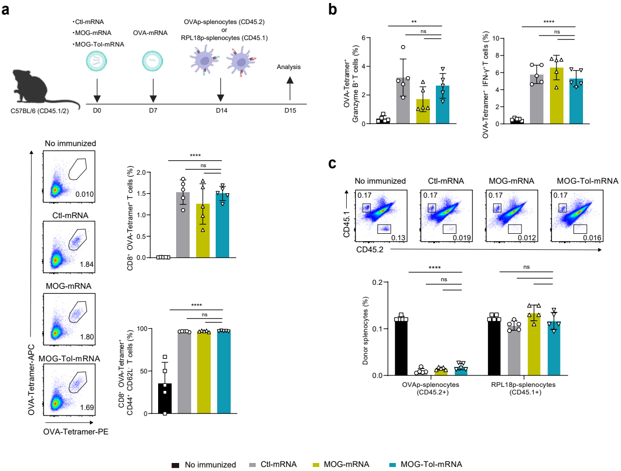


### Extended Data Figure 9

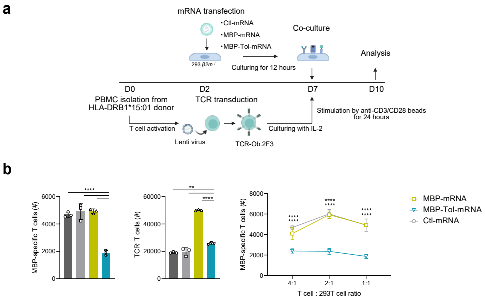


### Extended Data Figure 10

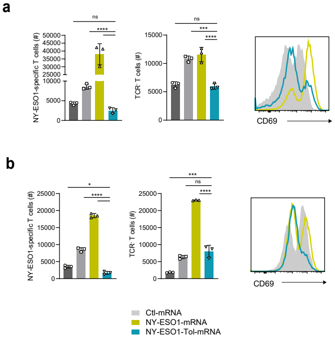


### Extended Data Figure 11
